# Cysteine dependent conformation heterogeneity of *Shigella flexneri* autotransporter IcsA and implications in its function

**DOI:** 10.1101/2022.07.21.501072

**Authors:** Jilong Qin, Yaoqin Hong, Renato Morona, Makrina Totsika

**Author notes:** Correspondence: Jilong Qin, Makrina Totsika:.

## Abstract

*Shigella* IcsA is a versatile surface virulence factor required for both early and late pathogenesis stages, extracellularly to intracellularly. Despite IcsA serving as a model Type V secretion system (T5SS) autotransporter to study host pathogen interactions, its detailed molecular architecture is poorly understood. Recently, IcsA was found to switch to a different conformation for its adhesin activity upon sensing of the host stimuli by *Shigella* Type III secretion system (T3SS). Here, we report that the single cysteine residue (C130) near the N-terminus of IcsA passenger has a role in IcsA adhesin activity. We also show that the IcsA passenger (IcsAp) exists in multiple conformations, and the conformation populations are influenced by a central pair of cysteine residues (C375 and C379), which is not previously reported for any Type V autotransporter passengers. Disruption of either or both central cysteine residues alters the exposure of IcsA epitopes to polyclonal anti-IcsA antibodies previously shown to block *Shigella* adherence, yet without loss of IcsA intracellular functions in actin-based motility (ABM). Anti-IcsA antibody reactivity was restored when the IcsA paired cysteine substitution mutants were expressed in a *∆ipaD* background with a constitutively active T3SS, highlighting an interplay between T3SS and T5SS. The work here uncovers a novel molecular switch empowered by a centrally localised, short-spaced cysteine pair in the Type V autotransporter IcsA that ensures conformational heterogeneity to aid IcsA evasion of host immunity.

**Importance:** *Shigella* species are the leading cause of diarrheal related death globally by causing bacillary dysentery. The surface virulence factor IcsA which is essential for *Shigella* pathogenesis is a unique multi-functional autotransporter that is responsible for cell adhesion, and actin-based motility, yet detailed mechanistic understanding is lacking. Here, we show that the three cysteine residues in IcsA contribute to the protein’s distinct functions. The N terminus cysteine residue within the IcsA passenger domain plays a role in adhesin function, while a centrally localised cysteine pair provides conformational heterogeneity resulting in IcsA molecules with different reactivity to adhesion-blocking anti-IcsA antibodies. In synergy with the Type III secretion system, this molecular switch preserves biological function in distinct IcsA conformations for cell adhesion, actin-based motility and autophagy escape, providing a potential strategy by which *Shigella* evade host immunity targeting of this essential virulence factor.

## Introduction

*Shigella* is a Gram-negative bacterial pathogen which is estimated to cause 80-165 million cases of shigellosis, with over 600,000 deaths worldwide annually (1). One of the essential *Shigella* virulence determinants (2) is the unipolarly distributed surface protein IcsA (formerly VirG) (3). In the human gut lumen, *Shigella* senses the bile salt deoxycholate (DOC) (4) via the Type III secretion system (T3SS) needle residing protein IpaD (5) to regulate IcsA’s adhesin activity required for pathogenesis (6). Inside the host cells, *Shigella* uses IcsA to perform actin-based motility (ABM) for inter- and intra-cellular spreading by interacting with host <u>N</u>eural-<u>W</u>iskott <u>A</u>ldrich <u>S</u>yndrome <u>p</u>rotein (N-WASP) (7). In addition, IcsA was also found to be a target of host autophagy recognised by ATG5, a critical host protein for the initiation of autophagosome formation (8). However, *Shigella* efficiently escapes host autophagy by masking the IcsA region targeted by ATG5 with the *Shigella* T3SS effector protein IcsB (8) (Figure 1a).

**Figure 1.**
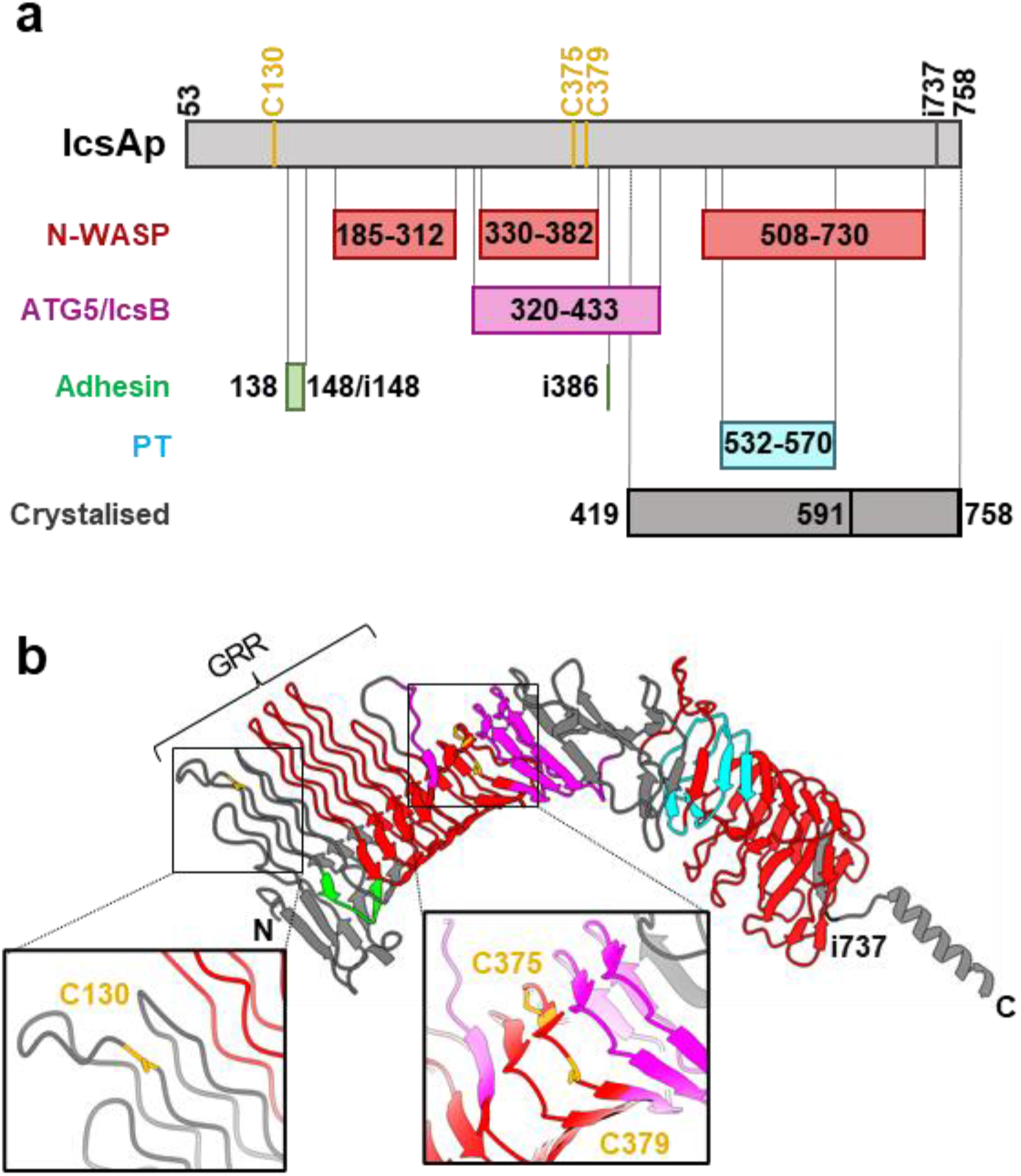
The autotransporter protein IcsA passenger domain. **a)** Schematic representation of defined subregions on IcsA passenger affecting its known biological functions with all three cysteine positions shown. PT, polar targeting. **b)** IcsA passenger tertiary structure predicted by AlphaFold (AF-Q7BCK4-F1) with defined functional subregions marked in colour as in **a)**, and cysteine residue locations show in close-up view. GRR, glycine-rich repeats (aa 117-307).

IcsA is encoded by the *Shigella* virulence plasmid (2) and is characterised as a type Va autotransporter (9) with typical protein domain architecture, including an N-terminal signal peptide (aa 1-52), a ∼75 kDa central passenger domain (aa 53-758) and an anti-paralleled beta-barrel (aa 759-1102) that is embedded in the outer membrane (OM) for the translocation of the IcsA passenger (IcsAp). Approximately 20% of IcsAp is cleaved off by the OM protease IcsP after translocation, with the remaining having a unipolar localisation on the bacterial surface (10). Protein subregions important for IcsA adhesin activity (6, 11), ABM function (12, 13), initiation of host autophagy (8), as well as the bacterial unipolar targeting (14) are all contained within the IcsAp domain (Figure 1a). However, the structure of IcsAp is only available for its C-terminal portions (591-758 and 419-758) and thus detailed mechanistic understanding of how multiple functions are ensured by this protein remains limited (15, 16) (Figure 1a).

The translocation of the passenger domain in type Va autotransporters is generally assumed to occur through the hairpin model, where passenger translocation proceeds from the carboxyl-terminal to amino-terminal through hairpin formation between a static strand sequestered in the lumen of the barrel and a sliding strand moving through the pore (17). This posits that the passenger remains unfolded in the periplasm, which has been supported by evidence whereupon a large polypeptide loop formed by an intramolecular disulfide bond prior to translocation inhibited the translocation of autotransporter passenger domain across the OM (18-21). This also aligns with the observation that most autotransporters have a low cysteine content in their passenger domain (22). However, non-autotransporter proteins containing a cysteine disulfide loop were shown to be successfully translocated when fused to autotransporter barrel domains (23, 24). A few autotransporters have a single cysteine pair in their passenger domain, with the most common spacing being 11 residues and proximal to the C terminal region of the passenger (25). Substitutions of such paired cysteine residues in the passenger domain of *Helicobacter pylori* VacA and *Serratia marcescens* Ssp-1 resulted in decreased protein production (25, 26). However, the role of cysteine residues in the passenger domain of type V autotransporters is poorly understood.

IcsA is an unique autotransporter and is distinctive to other cysteine containing autotransporters in that its paired cysteines are very short spaced (4 aa between C375 and C379) and located at the central region of the passenger, as well as the presence of an additional unpaired cysteine residue (C130) at the N-terminal region of IcsAp (Figure 1a) (22). IcsA was reported previously to form an intramolecular disulfide bond in the periplasm prior to OM translocation in a DsbB dependent manner (27). However, the exact cysteine residues participating in the intramolecular disulfide bond were not defined. In the AlphaFold predicted IcsA structure (Figure 1b) (28), the unpaired C130 is located at the glycine-rich repeats (GRR) region close to the previously identified adhesin region (11) (Figure 1a). The central paired cysteine residues (C375 and C379) are located at the beta-helical backbone near the ∼90° kink region (16), which overlaps with both the N-WASP interacting region II (IRII) and the ATG5/IcsB binding region, and are close to the other potential adhesin site (i386) identified previously (6, 8, 12) (Figure 1a). However, no intramolecular disulfide bond is represented in the predicted structure and whether these cysteines are important for IcsA biogenesis and/or biological functions remains unknown. Here, we investigated the role of the three cysteines in IcsAp biogenesis, conformation and function. We showed that the unpaired cysteine residue C130 affects IcsA’s adhesin activity, and that the paired cysteine residues form an intramolecular disulfide bond that impacts on IcsA conformational heterogeneity.

## Results

### Cysteine substitution IcsA passenger mutants have altered conformation

We investigated the role of cysteine residues (C130, C375 and C379) within IcsAp in IcsA biogenesis and biological functions. We substituted each cysteine residue individually (C130S, C375S and C379S) or in a pair (C375S/C379S, double mutant, DM) to serine to minimise potential structural perturbations. All substitutions did not affect IcsA expression levels (Figure 2a). Interestingly, substitution of the paired cysteine residues (C375 and C379) either individually or in combination, but not of unpaired cysteine residue (C130), abolished IcsA detection on the cell surface by indirect surface immunofluorescent labelling with anti-IcsA polyclonal antibodies (Figure 2b) which had previously been shown to neutralise *Shigella* adherence to host cells (11). We reasoned that loss of detection of the paired cysteine substitution mutants could be due to either a translocation defect, or altered accessibility of epitopes recognised by the anti-IcsA antibodies used. To distinguish between the two hypotheses, we performed proteinase surface shaving of intact *S. flexneri* bacteria expressing mutant IcsA proteins. We found that all IcsA cysteine substitution mutants remain sensitive to extracellular proteinase K digestion similar to WT (Figure 2c), suggesting that the passenger domain of IcsA is successfully translocated to the surface. Indeed, when we precipitated the secreted extracellular protein from culture supernatants, we found all IcsA cysteine substitution mutants secreted IcsAp in a similar level to IcsA^WT^ (Figure 2d), confirming that passenger translocation was not affected. Thus, the loss of surface detection of the IcsA paired cysteine substitution mutants by anti-IcsA antibodies was most likely due to altered accessibility of epitopes to anti-IcsA antibodies. To investigate this, and examine possible polar targeting defects, we engineered a FLAG×3 tag at the C terminus of IcsAp (i737, Figure 1) and confirmed no impact on IcsA production (Figure 2e). We then performed surface immunofluorescent labelling with both anti-IcsA and anti-FLAG antibodies. The anti-FLAG antibody, but not anti-IcsA antibodies, successfully labelled all IcsA cysteine substitution mutants and confirmed surface localisation of IcsAp from the paired cysteine mutants (Figure 2f). Image analysis confirmed that none of the cysteine substitution mutants have defects in polar targeting (Figure 2f). Taken together, these data indicate that paired cysteine residues in IcsAp impact IcsA conformation, but not translocation, surface localisation or extracellular cleavage.

**Figure 2.**
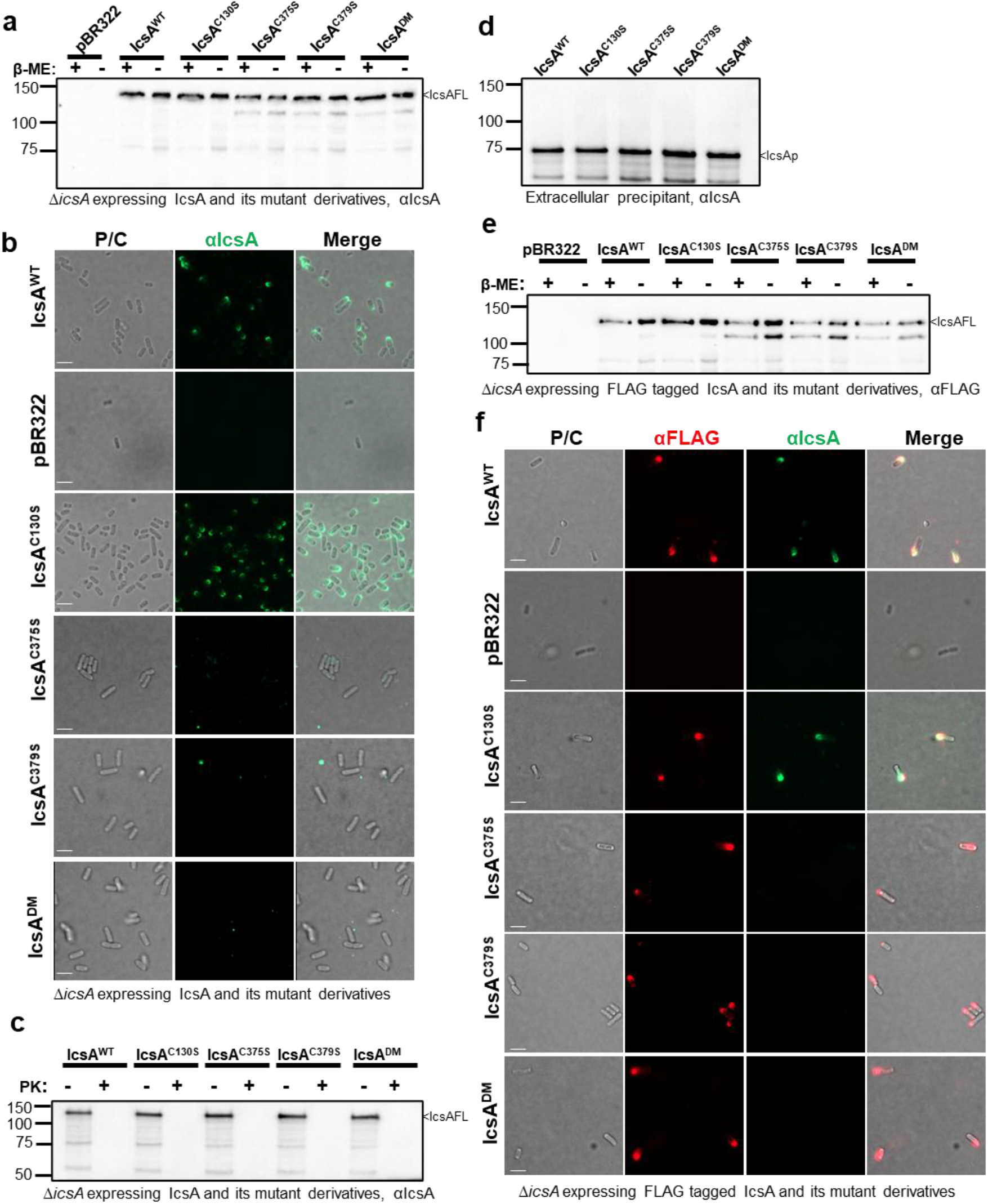
Cysteine substitutions in IcsAp do not affect surface display but influence protein conformation. **a)** Western immunoblotting of bacterial cell lysates expressing IcsA^WT^ and cysteine mutant detected with anti-IcsA antibodies. IcsAFL, IcsA full length. **b)** Indirect bacterial surface immunofluorescent labelling of *S. flexneri* expressing IcsA^WT^ and cysteine substitution mutants with anti-IcsA antibodies. **c)** Western immunoblotting of IcsA with lysates of bacterial cells with and without proteinase K surface shaving. **d)** Western immunoblotting of precipitated extracellular cleaved IcsAp. **e)** Western immunoblotting of lysates of bacterial cells expressed FLAG-tagged IcsA^WT^ and cysteine substitution mutants with anti-FLAG antibodies. **f)** Indirect bacterial surface immunofluorescent labelling of FLAG-tagged IcsA passenger with anti-IcsA antibodies and anti-FLAG antibody. Scale bars shown as 2 μm in **b)** and **f)**.

### Paired Cys substitution in IcsA alters its N-terminus conformation and the conformational difference is maintained after secretion

To characterise the IcsAp conformational differences due to cysteine substitution mutations on the bacterial surface, we performed limited proteolysis with human neutrophil elastase (hNE) to surface shave IcsAp in a pulse-chase manner. Under immediate exposure of hNE, the IcsA^WT^ and IcsA^C130S^ were digested in a ∼100 kDa fragment, followed by a ∼80 kDa fragment (Figure 3a, red arrow) at 5 min, and a ∼40 kDa fragment for the rest of 85 min digestion duration (Figure 3a, black arrow). These observations suggested that the hNE accessibility was not detectably altered between IcsA^WT^ and IcsA^C130S^. In contrast, all paired cysteine substitution mutants were more resistant immediately after exposure to hNE, producing a fragment running higher than that of IcsA^WT^ (∼110 kDa) (Figure 3a, blue arrow), followed by a ∼90 kDa fragment at 5 min (Figure 3a, green arrow). The ∼40 kDa band (Figure 3a, black arrow) appearing after 5 min digestion was consistently detected for all IcsA mutants and IcsA^WT^.

**Figure 3.**
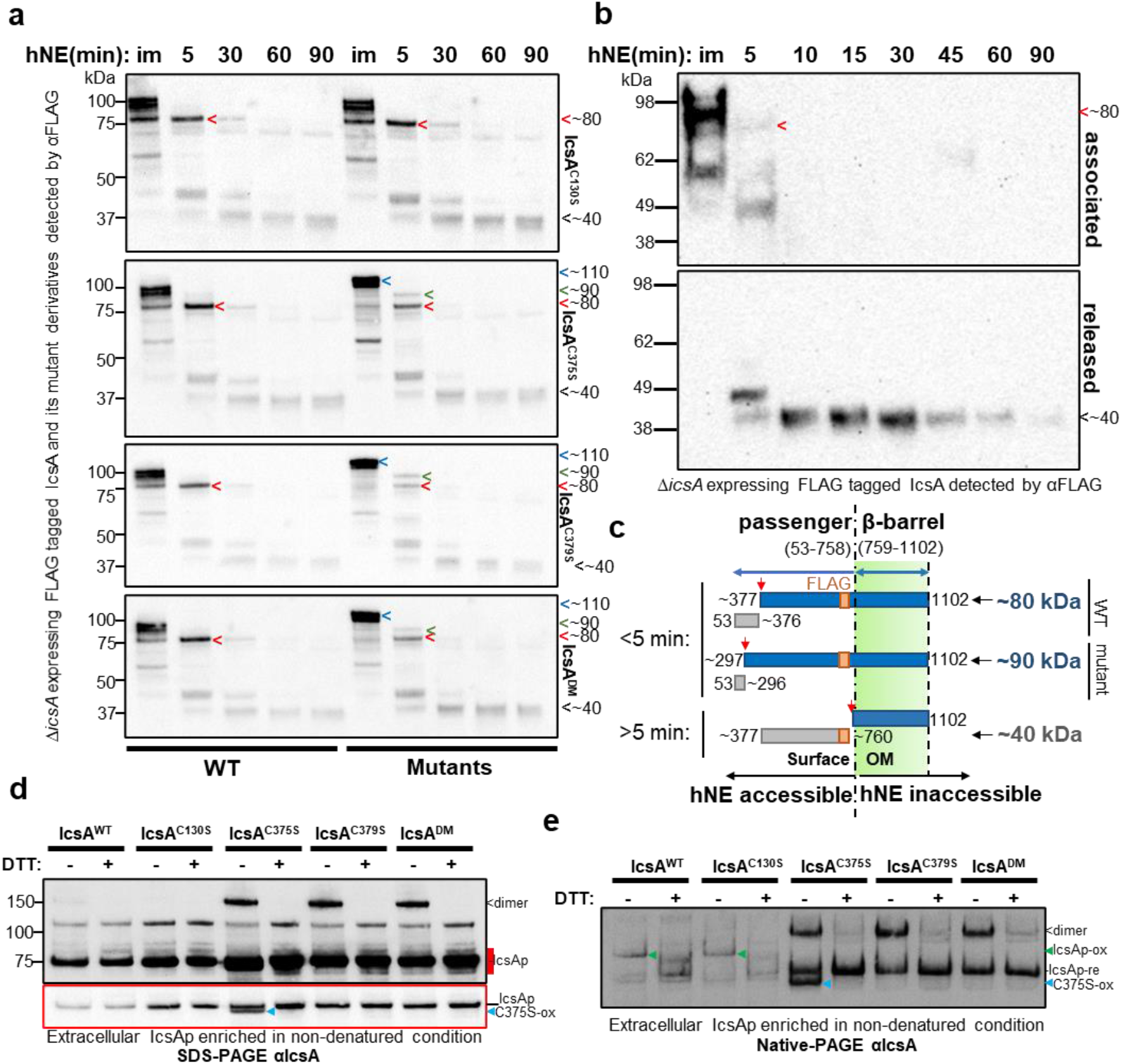
The IcsA N terminus has the altered conformation and was maintained in the cleaved IcsAp. **a)** Western immunoblotting with bacterial cells expressing IcsA and its cysteine substitution mutants treated with hNE. Samples were taken at different time points as indicated above the blot. im, immediate recovery of the digestion sample upon the addition of hNE. Samples from bacteria expressing IcsA^WT^ were electrophoresed in parallel (left) with mutants (right) and blotted together to ensure equal exposure for direct comparison. **b)** Western immunoblotting of fractionated hNE digestion samples from the bacteria expressing IcsA^WT^. **c)** Model of the digestion differences due to altered conformation in IcsA mutant with estimated cutting sites shown. **d)** Western immunoblotting of IcsAp separated in semi-denatured conditions. Extracellular IcsA was precipitated in ammonium sulfate and was solubilised in SDS-PAGE buffer in the presence or absence of DTT, and was subjected to SDS-PAGE and immunoblotting without heating. The under-exposure image in red border correlates to the region in the longer exposure image was marked red. re, reduced; ox, oxidised. <: IcsAp dimer; blue arrow: oxidised IcsA^C375S^, C375S-ox. **e)** Culture supernatant IcsAp protein recovered as above analysed by Native-PAGE and detected with anti-IcsA antibodies. green arrow: oxidised IcsAp, IcsAp-ox.

To further distinguish the cell association states of these FLAG-tagged IcsA fragments, we cell fractionated the digestion reaction for each time point, and blotting revealed that the ∼40 kDa FLAG-tagged fragment is cleaved off the bacterial cell surface (Figure 3b). Hence, it can be deduced that the FLAG-tagged IcsA fragment at ∼40 kDa is devoid of the beta-barrel domain (inaccessible to hNE) as it is not cell associated. HNe was reported previously to preferably cleave valine, alanine, and isoleucine (29). Therefore, it can be estimated that the ∼40 kDa fragment is derived from aa 377-760 (FLAG engineered at i737) (Figure 3c), suggesting that the accessibility to hNE in this region is unchanged for all cysteine substitution mutants. In contrast, the larger FLAG-tagged IcsA ∼80 kDa fragment, which is slightly larger than the predicted size of full length IcsAp (∼75 kDa), remains cell associated (Figure 3b, red arrow) and therefore is uncleaved from the beta-barrel. Hence it can be estimated to be derived from aa 377-1102 (Figure 3c). Similarly, the ∼90 kDa FLAG-tagged fragment detected from the paired cysteine substitution mutant IcsA can be estimated to be derived from aa 297-1102 (Figure 3c). These data indicated that the altered hNE accessibility of IcsAp for all paired cysteine substitution mutants are at the N-terminus region estimated between aa 53-376, which is the N-terminus side of C375 and C379. This prediction is also in agreement with the differential surface immunostaining observed with the anti-IcsA antibodies previously (Figure 2 b&f), which were confirmed to only react with IcsAp (30).

IcsAp can be cleaved off the bacterial surface by IcsP and be secreted into the extracellular medium (10). To rule out the possibility that the observed changes for the paired cysteine substitution mutants were due to differential LPS masking, we compared all secreted IcsA cysteine substitution mutant passengers to that of the IcsA^WT^ in semi-denatured conditions in the presence or absence of the disulfide bond reducing agent DTT (Figure 3d). As expected, following recovery from the culture supernatant, all cleaved IcsAp were detected at the molecular size of ∼75 kDa (Figure 3d, IcsAp). Interestingly, a DTT sensitive IcsAp population with a molecular size that correlates to an IcsA dimer (150 kDa, Figure 3d, dimer) was detected with IcsA^WT^ and all paired cysteine substitution mutants but not with the IcsAp^C130S^, suggesting that C130 is involved in an intermolecular disulfide bond formed between two IcsAp molecules under our experimental conditions. We also detected a faster migrating and DTT sensitive IcsAp band only with the IcsA^C375S^ mutant (Figure 3d, C375S-ox). Presumably, this was due to the oxidation of the two remaining cysteines C130 and C379, leading to a slightly more compact molecule under semi-denatured conditions. This also suggests that C130 is located close to C379 in the folded IcsAp structure.

Next, to capture differences in protein conformation mediated by the paired cysteines that could not be potentially resolved under semi-denatured conditions due to the small size of the disulfide loop formed by C375-C379, we compared all cysteine substitution mutants in non-denatured gel conditions (Figure 3e). Under these conditions, migration speed changes potentially for proteins in different conformations due to changes in molecular shape, oligomeric state, and exposure of their intrinsic surface charge. Under non-denaturing conditions, we still detected the potential dimeric form of IcsAp, that was predominantly enriched in the paired cysteine substitution mutants (Figure 3e, dimer) and the oxidised IcsAp^C375S^ (Figure 3e, C375S-ox). In addition, we detected a DTT resistant IcsAp for IcsAp^WT^ and all cysteine substitution mutants (Figure 3e, IcsAp-re). In contrast, IcsAp^WT^ and the IcsAp^C130S^ were instead detected with an additional DTT sensitive IcsAp form migrating at a different speed (Figure 3e, IcsAp-ox) relative to those detected with paired cysteine mutant IcsAp. It is likely that the populations (Figure 3e, IcsAp-ox) observed for IcsAp^WT^ and IcsAp^C130S^ were due to intramolecular oxidation between C375 and C379, since this population is absent in the paired cysteine substitution mutants (Figure 3e, IcsAp-re). Together, these data suggest IcsA conformation is maintained through a disulfide bond formed between C375 and C379, and that the IcsAp conformation is maintained even after IcsP cleavage.

### Purified IcsA exists in two conformations

To show that the conformations of IcsAp are independent of binding to other secreted *S. flenxeri* effector proteins, we adapted a similar expression and purification strategy to that used previously (31) to isolate native N-terminus FLAG tagged IcsAp (Figure 4a) from the culture supernatant. Same as reported previously, we separated IcsAp into two fractions through size exclusion chromatography (SEC) (Figure 4b&c) (31) with the latter fraction (migrating as 137 kDa determined with native molecular weight markers with globular shape) correlated to a monomer determined by SEC coupled with multiangle light scattering (SEC-MALS) in the previous study (31). These two IcsAp species can also be separated using Native-PAGE (Figure 4d) with the migrating molecular size correlated to SEC results (Figure 4b). We are aware that the protein molecular weight determination with the globular molecular weight markers is irrelevant since the shape of IcsAp is predicted to be elongated, and hence we are only using them as a relative mass reference to correlate our SEC and Native PAGE results.

**Figure 4.**
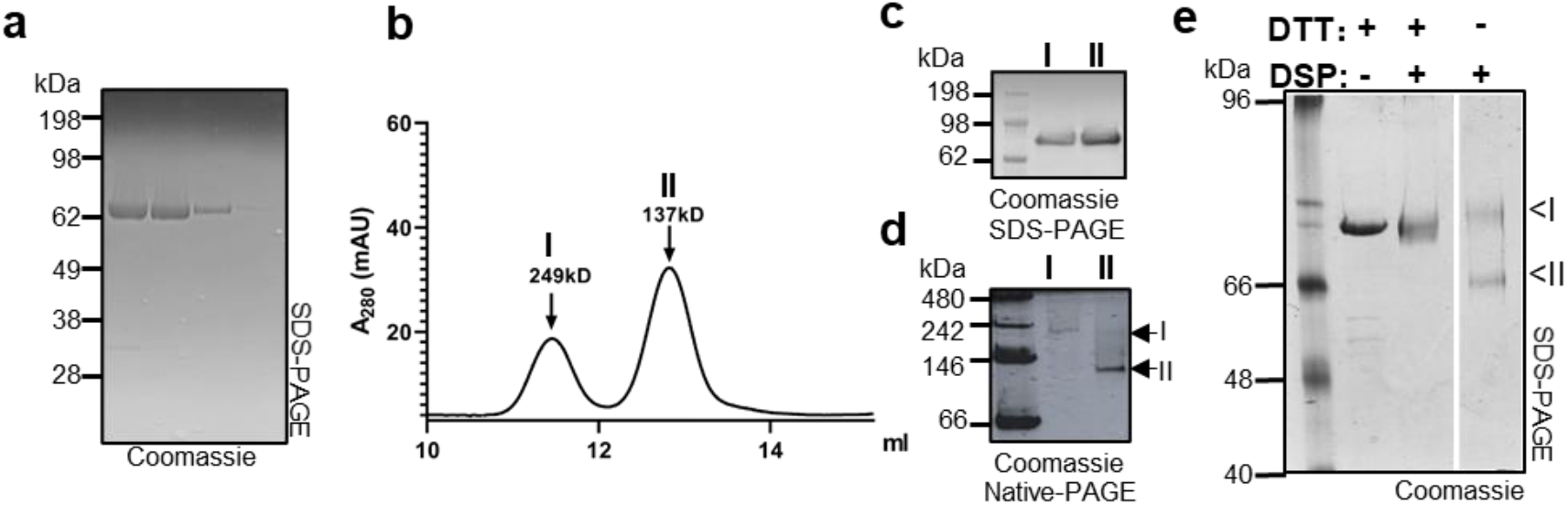
Purified IcsA exists in two different conformations. **a)** Affinity purified IcsAp resolved by SDS-PAGE and stained with Coomassie blue. **b)** SEC separation of IcsAp into populations I and II. **c)** SDS-PAGE of two IcsAp populations isolated by SEC. **d)** Native-PAGE of two IcsAp populations isolated by SEC. **e)** SDS-PAGE of IcsAp crosslinked with 0.1 mM DSP in the presence or absence of 10 mM DTT. Image is from the same gel with relevant area shown.

To prove that purified IcsAp has two forms adopting different protein conformations, we attempted to fix any close intramolecular contacts by incubating purified IcsAp protein with the DTT reducible crosslinker DSP (Figure 4a) and comparing the migration profile under denaturing gel conditions. Upon crosslinking with DSP, we resolved an IcsAp population with a faster migration (Figure 4e, II), suggesting a compact conformation and a closer intramolecular contact than the alternative IcsAp conformation (Figure 4e, I). Together, these data suggest that the IcsAp exists in multiple conformations with a compact form that allows closer intramolecular contacts, and an extended form with fewer intramolecular contacts.

### Effects of IcsA cysteine substitution mutations in Shigella host cell adherence and infection

IcsA is well known as a multifunctional virulence factor with defined regions interacting with different host molecules (Figure 1a) facilitating *Shigella* pathogenesis at different stages. Here we established that the cysteine substitutions in IcsA passenger domain altered IcsA oligomeric states and conformations, and even altered the accessibility of polyclonal anti-IcsA antibodies. We then investigated whether these altered conformations due to cysteine substitution would also alter the interactions of IcsA with host molecules, thereby having functional implications. IcsA was reported previously to act as an adhesin interacting with host cell surfaces when T3SS is activated (6). We therefore expressed all IcsA cysteine substitution mutants in a *S. flexneri ∆ipaD* strain background, where the loss of IpaD results in hyper-adhesion, similar to IpaD binding to the host environmental stimulus deoxycholate (DOC). The strain producing IcsA^C130S^, but not strains producing paired cysteine substitution IcsA mutants (IcsA^C375S^, IcsA^C379S^ and IcsA^DM^), lost hyper-adhesion to host cells in comparison to controls with IcsA^WT^ (Figure 5a). However, none of the strains producing IcsA mutants had a defect in cell invasion (Figure 5b). In addition, although the paired cysteine residues are in the N-WASP protein interaction region II (Figure 1a) which was previously reported to be important for ABM function (12), we observed no differences in intracellular F-actin polymerisation (Figure 5c) for the strains with IcsA cysteine substitution mutants compared to WT. This is also supported by the plaque formation assay which showed no defects in intercellular spreading ability owing to the normal ABM function for strains with IcsA mutants (Figure 5d & Figure S1a). The region where C375 and C379 are located (Figure 1a) was also reported to be a target of host autophagy through the recognition and binding of the host autophagosome component ATG5, yet intracellular *Shigella* can escape this host defence mechanism by masking the same region on IcsA by binding to IcsB (8). Indeed, *S. flexneri* lacking IcsB produced smaller plaques and this could be restored by *in trans* complementation of IcsB and its fusion chaperone complexes (Figure 5e & Figure S1b) as reported before (32). We then examined if the altered conformation in IcsAp seen with the paired cysteine mutants would be naturally resistant to host autophagy without IcsB masking due to any potentially altered accessibility to host ATG5. We did not detect any restored plaque sizes for any strains with IcsA cysteine substitution mutants in the absence of IcsB masking (Figure 5e & Figure S1b). Taken together, our data suggest that the C130 residue of IcsA is involved in adhesin function consistent with its location close to the previously defined adhesion region (11), while the altered conformations due to the paired cysteine substitutions are tolerant to ABM function and escape from autophagy under our experimental conditions.

**Figure 5.**
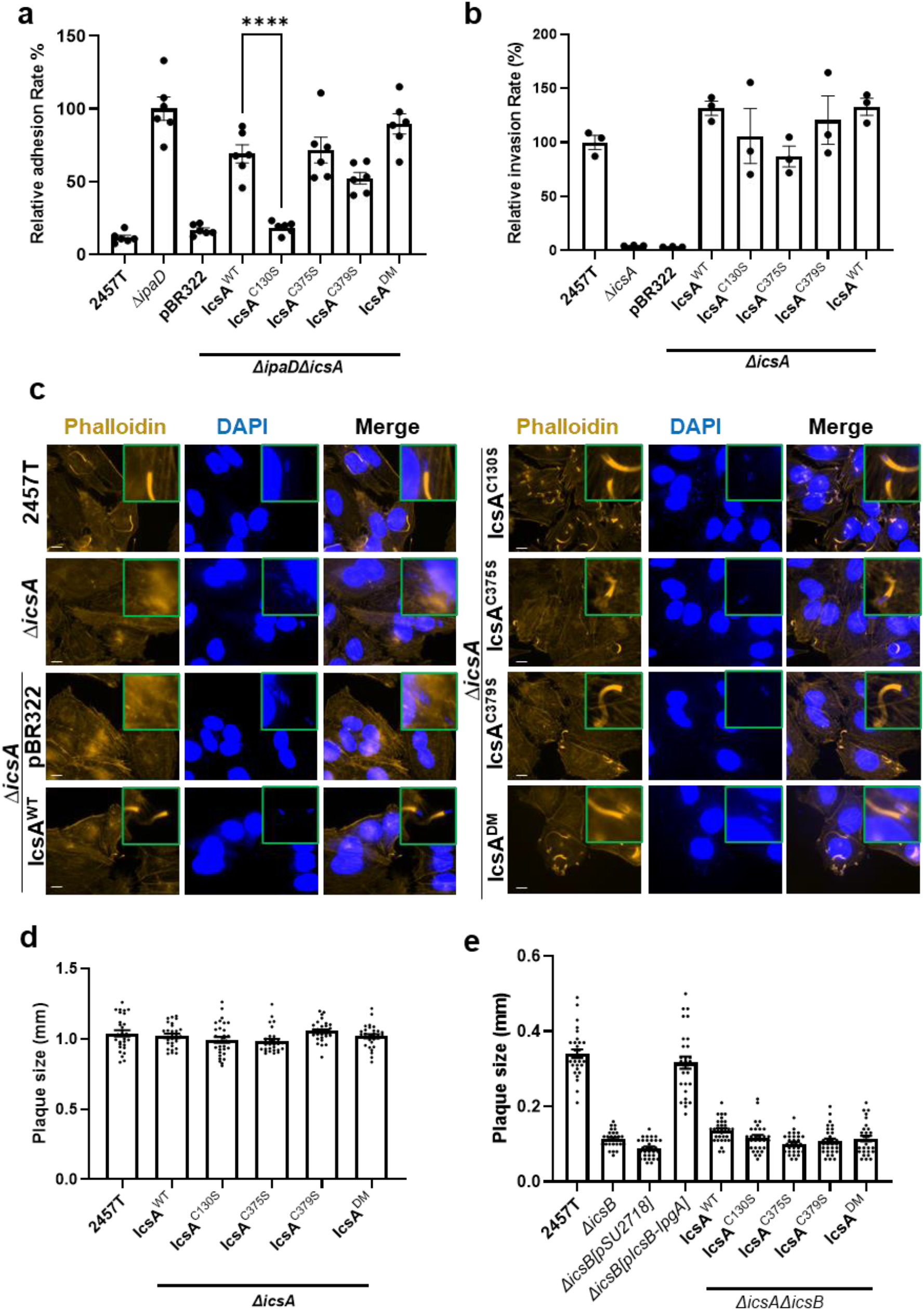
Effect of cysteine substitution mutation on IcsA function. **a)** Cell adhesion assay of indicated *S. flexneri* strains with HeLa cells. Mean values from 6 independent assays with standard error mean (SEM) are shown. Statistical significance was calculated using one-way ANOVA followed by Dunnett’s multiple comparisons test against *∆ipaD∆icsA* pIcsA^WT^, and p values are as follows: ****, p<0.0001. **b)** HeLa cell invasion assay of indicated *S. flexneri* strains treated with 2.5 mM DOC. Data represents 3 independent assays. **c)** Fluorescent labelling of host cell F-actin by phalloidin and host and bacterial nuclei by DAPI of HeLa cells infected with indicated *S. flexneri* strains. Scale bar as 10 μm. **d)** and **e)** Plaque sizes formed by indicated *S. flexneri* strains in MDCK-2 monolayers. Data were from two independent assays with ∼30 plaque sizes plotted for each strain.

### The ΔipaD mutation restores anti-IcsA accessibility by further alterations in IcsA conformation

The IcsA paired cysteine substitution mutants (IcsA^C375S^, IcsA^C379S^ and IcsA^DM^) exhibited dramatic conformational changes that masked epitopes reactive with the anti-IcsA antibodies used, yet the proteins were still able to polymerises F-actin and be the target of host autophagy when exposed intracellularly. It is known that the *Shigella* type III secretion system (T3SS) can be activated through the binding of IpaD to DOC, which in turn alters IcsA conformations (6). Potentially such conformational changes may rescue the altered accessibility to anti-IcsA antibodies in the paired cysteine substitution mutants. Indeed, when expressed in a *S. flexneri ∆ipaD∆icsA* background with constitutively active T3SS, the surface detection of IcsA^C375S^, IcsA^C379S^ and IcsA^DM^ by anti-IcsA antibodies was restored (Figure 6a). We then investigated the conformation of secreted IcsAp of all the IcsA mutants in both semi-denatured (Figure 6b) and nondenatured conditions (Figure 6c). We consistently detected the IcsAp population with the molecular mass correlated to a dimerised IcsA complex (in an C130 dependent manner) (Figure 6b, dimer), as well as the oxidised IcsAp^C375S^ population only with the strain expressing IcsA^C375S^ (Figure 6b, C375S-ox). However, the amount of the secreted IcsAp dimer population (Figure 6 b&c, dimer) for the paired cysteine substitution IcsA mutants was decreased when expressed in the hyper-adhesive strain *S. flexneri ∆icsA∆ipaD* in comparison to *S. flexneri ∆icsA* (Figure 6c, dimer, and Table 1). In contrast, the level of the other conformation population (Figure 6c, left, IcsAp-re) for these paired cysteine substitution IcsA mutants was increased and was resistant to DTT treatment (Figure 6c, right, IcsAp-re). A similar conformation population increase in *S. flexneri ∆icsA∆ipaD* was also observed for IcsAp^WT^ and IcsAp^C130S^, albeit in an oxidised form (Figure 6c, left, IcsAp-ox). The oxidised form detected in IcsAp^WT^ and IcsAp^C130S^ is potentially mediated by a disulfide bond between C375 and C379 (Figure 6c, left, IcsAp-ox), since they migrate the same as the paired cysteine substitution mutant IcsAp upon DTT reduction (Figure 6c, right, IcsAp-re). Collectively, IcsAp expressed from *S. flexneri ∆icsA∆ipaD* favours a conformation with less intermolecular interactions with a decrease in the C130-mediated IcsA dimer population. These data suggest that the activation of T3SS alters the conformation landscape of IcsA to restore its epitope exposure to anti-IcsA antibodies used.

**Table 1.** Cysteine-dependent IcsA multi-conformations and their functional implications.

|  |  | *S. flexneri ΔicsA* |  |  |  |  | *S. flexneri ΔicsAΔipaD* |  |  |  |  |
| --- | --- | --- | --- | --- | --- | --- | --- | --- | --- | --- | --- |
| IcsA type |  | WT | C130S | C375S | C379S | DM | WT | C130S | C375S | C379S | DM |
| anti-IcsA reactivity |  | + | + | - | - | - | + | + | + | + | + |
| ABM |  | + | + | + | + | + | nt | nt | nt | nt | nt |
| Adhesion |  | nt | nt | nt | nt | nt | + | - | + | + | + |
| Invasion |  | + | + | + | + | + | nt | nt | nt | nt | nt |
| AE with *ΔicsB* |  | - | - | - | - | - | nt | nt | nt | nt | nt |
| IcsAp conformations | IcsAp-re | low | low | high | high | high | low | low | high | high | high |
|  | IcsAp-ox | high | high | nd | nd | nd | high | high | nd | nd | nd |
|  | Dimer | low | nd | high | high | high | low | nd | low | low | low |
|  | C375S-ox | low | low | high | nd | nd | low | low | high | nd | nd |

The relative level of IcsAp conformation populations detected by Western immunoblotting were reported. AE, autophagy escape; DM, C375S/C379S double mutant; nt, not tested; nd, nondetectable.

**Figure 6.**
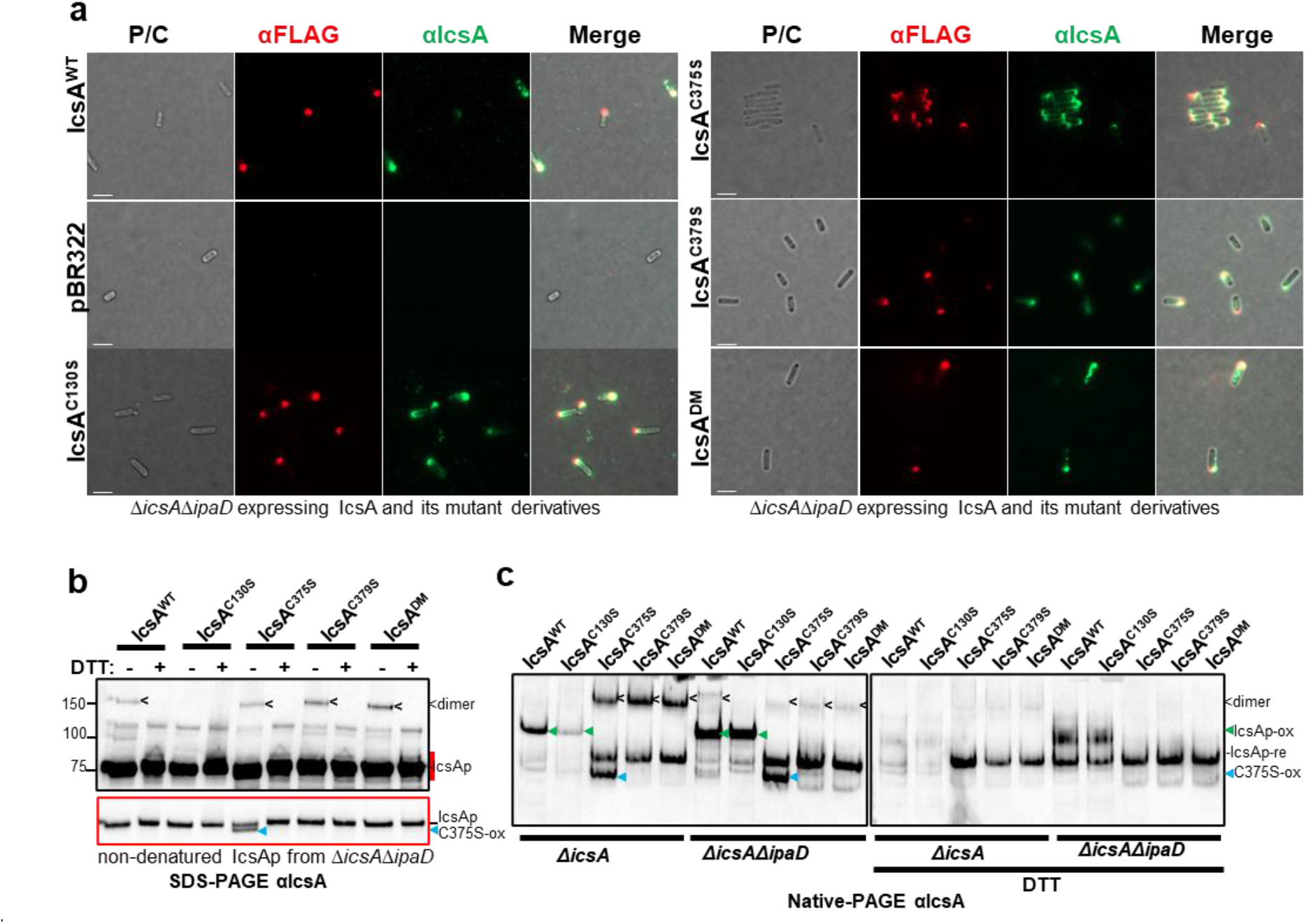
Effect of *ΔipaD* on IcsA conformation. **a)** Indirect bacterial surface immunofluorescent labelling of FLAG-tagged IcsA passenger with anti-IcsA antibodies and anti-FLAG antibody. **b)** Western immunoblotting of IcsAp recovered from culture supernatant of indicated *S. flexneri* strains expressing IcsA and IcsA mutants in non-denatured conditions. Samples were solubilised in SDS-PAGE buffer with or without DTT, and subjected to SDS-PAGE and Western immunoblotting without heating. The under-exposure image in red border correlates to the region in the longer exposure image was marked red. re: reduced, ox: oxidised, <: IcsAp dimer; blue arrow: oxidised IcsA^C375S^, C375S-ox. **c)** IcsAp protein recovered as above and analysed by Native-PAGE and detected with anti-IcsA antibodies. green arrow: oxidised IcsAp, IcsAp-ox.

## Discussion

IcsA is a multi-functional virulence factor being strictly required for different stages of *Shigella* pathogenesis. The configuration of bearing a central cysteine pair and a N-terminal unpaired cysteine residue makes IcsA an unique type Va autotransporter. We here studied the role of these three cysteine residues on IcsA’s structure and biological functions (Table 1).

We discovered that the unpaired cysteine residue C130 is involved in IcsA’s adhesin activity. C130 is located close to the previously identified region (138-148) affecting IcsA adhesin function (11). Deletion of either this entire region or insertion by a linker (5aa) at Q148 affected IcsA adhesin activity (6, 11). It is possible that the unpaired cysteine residue C130 is directly contributing to the host cell adherence, as the disruption of the upstream folding at the C-terminus would then affect the subsequent folding of this region during the later stage of OM translocation and may orient C130 differently. In support of this speculation, substitution Q148C also completely abolished IcsA’s adhesin activity, possibly due to disulfide bond formation between C148 and C130 (11). A single cysteine residue has also been reported previously to be responsible for the activity of adhesin Ifp in *Yersinia pseudotuberculosis* (33).

We also detected an IcsA dimer population that is formed by a disulfide bond between C130 residues from two different protein molecules. It is unlikely that the intermolecular disulfide bond was formed in the periplasm, and instead is likely to be due to post-translocation oxidation by the environment. IcsA has been reported previously to have self-association activity (34), and mutants were demonstrated to restore the IcsA N-WASP binding defect cooperatively (13), yet the exact region for the self-association remains unclear. It is possible that such self-association orients the two C130 residues from two IcsA molecules to form a disulfide bond.

By substituting the paired cysteine residues with serine, either individually or together, we revealed that the centrally localised cysteine pair (C375/C379) form an intramolecular disulfide bond. IcsA has been reported previously to possess an intramolecular disulfide bond catalysed by the periplasmic disulfide bond formation enzymes DsbB/DsbA prior to OM translocation (27). Here we have shown that the intramolecular disulfide bond in IcsAp is maintained after the OM translocation and its subsequent release into extracellular milieu by IcsP. It has been shown previously with *H. pylori* VacA (25) and *S. marcescens* Ssp-1 (26) that disruption of intrinsic paired cysteine residues led to decreased protein production. However, we have shown here that disruption of paired cysteine residues in IcsAp does not influence protein production. We reasoned that this is due to both the localisation of and the spacing between the paired cysteine residues. Both VacA and Spp-1 have a pair of cysteine residues localised to the C-terminus of the passenger domain (25), which potentially is required to stabilise the initial translocation hairpins for efficient translocation of the whole passenger. In contrast, the central short-distanced cysteine pairs in IcsA are likely to be tolerated during later translocation events where the C-terminal beta-helical stable core (15) has been completely translocated. This is also shown in other autotransporters with engineered short-spaced cysteine pairs, where small cysteine loops are tolerated during the OM translocation of the autotransporter passengers (18, 21). However, here we discovered that these short-spaced cysteine pairs in the IcsAp instead influence the autotransporter conformation when displayed on the cell surface, and may act as a molecular constrain to influence the matured autotransporter conformations after translocation. This may be similar to those adjacent cysteine residues in non-autotransporters which were shown previously as a redox switch to allosterically influence protein conformations and function (35, 36). By comparing the differential exposure of proteolysis sites to hNE for all IcsA cysteine substitution mutants, we refined the region with altered conformation in those paired cysteine mutants to be limited to aa 52-376, which is at the N-terminus side rather than the C-terminus side of C375/C379. This is probably due to the order of passenger translocation, where C terminal beta-helical core is firstly folded to guide the subsequent folding of N-terminal regions. Disruption of the disulfide bond formation between C375 and C379 is likely to alter the local folding and influences the subsequent N-terminus protein conformations only. Our findings may also be a useful guide for the design of short-spaced cysteine pairs in autotransporter passengers to study their protein conformations, functions, and generalised folding mechanisms.

In addition, we have shown in detail that one of the purified IcsAp conformation populations has a close intramolecular contact which upon cross-linking fixed it into a compact molecule in denatured conditions, distinctive to the other conformational population. We speculate that this intramolecular interaction may occur between the central region and the N-terminal region of the IcsAp through a potential bridging loop as proposed previously (31), although it was not predicted by the Alphafold (Figure 1b) (28), which instead predicted IcsA is having no cysteine residues forming an intramolecular disulfide bond. The former is however, supported by our experimental data. Specifically, upon disruption of C375, C130 likely forms a disulfide bond with C379, suggesting an intramolecular contact which oriented these two residues from N-terminal and central passenger closely in space. This result may also partially explain the previous report where a linker insertion at position i386 also affected IcsA adhesin activity (6), as they might have a close contact with N-terminal regions.

IcsA holds great vaccine candidate potential, as it was demonstrated previously that the level of antibodies to IcsA significantly reduces the risks of shigellosis (37). High levels of anti-IcsA antibodies were also found as one major maternal antibody transferred to infant and may likely contributing to disease prevention during the first months of life (38). The correlated protection is likely to be through the prevention of early *Shigella* host cell attachment as was demonstrated previously (11) and not through antibody mediated complement killing (38). However, here we show that the binding of IcsAp to the adhesion blocking polyclonal anti-IcsA antibodies can be abolished upon the disruption of the centrally localised cysteine pair in IcsAp that is involved in intramolecular disulfide bond formation. IcsA exists in multiple conformations. Indeed, it has been reported previously that a monoclonal anti-IcsA antibody targeting the GRR region only stained 40% of the *Shigella* bacterial cell surfaces (39). We have also shown that the IcsAp region with conformational heterogeneity is estimated to be aa 53-376, overlapping with the entire GRR region. More importantly, this mAb work suggests that the differences in IcsA conformations are probably segregated at the single cell level. Here we have shown that IcsA’s population landscape is influenced by an intramolecular disulfide bond formed between the central cysteine pair, and disruption of the disulfide bond by substitution mutagenesis shifts the IcsA population into a conformation with no reactivity to the adhesion blocking antibodies (Table 1). However, in this conformation, we could not detect any defects in their cell infection function including cell adhesion, invasion and intracellular movement. It is possible that the cellular function of IcsA upon paired cysteine disruption might be also restored due to the further conformational changes such as upon the activation of T3SS as shown here, in a similar way to the restoration of antibody detection. Nevertheless, this may be a strategy whereby *Shigella* IcsA evades host immunity by having multiple conformations with different reactivity to antibodies elicited by the host, working closely with the T3SS to preserve its biological functions.

## Materials and Methods

### Bacterial strains, plasmids, and mammalian cell line

The bacterial strains and plasmids used in this work are listed in Table S1. Single colonies of *E. coli* bacterial strains grown overnight on Lennox Broth (LB) (40) agar (1.5% w/v) plates or single red *S. flexneri* colonies grown on Trypticase Soy Agar plates containing 0.03% (w/v) Congo Red were picked and grown overnight in LB at 37 °C for all experiments. Where appropriate, media were supplemented with ampicillin (Amp, 100 µg/ml), kanamycin (Kan, 50 µg/ml), streptomycin (Strep, 100 µg/ml) or chloramphenicol (Chl, 25 µg/ml). HeLa cells were routinely cultured and maintained with MEM (Gibco) supplemented with 5% (v/v) FBS, and MDCK-2 cells were cultured and maintained with MEM(Gibco) or DMEM (Gibco) supplemented with 5% (v/v) FBS (Gibco) at 37 °C with 5% CO_2_.

### Mutagenesis by allelic exchange and inverse PCR

*S. flenxeri ΔicsB and ΔicsAΔicsB* mutants were generated by Lambda Red mutagenesis as described previously (41) with the oligos listed in Table S1.

### Plasmid construction

For the IcsB expression construct, the coding sequence of IcsB and its chaperone protein IpgA along with their native promoter region was PCR amplified (Table S1) and restriction cloned into pSU2718 with XbaI and KpnI, resulting in pIcsB-IpgA.

For IcsA site-directed mutagenesis, pIcsA (42) was used as the template to amplify the entire plasmid by inverse PCR using oligos (Table S1) with cysteine to serine amino acid substitutions at the 5’ end. Amplicons were then phosphorylated, ligated, and introduced into TOP10 for recovery. Plasmids were then extracted from transformants, and the codon substitution was confirmed by sequencing. The in-frame addition of FLAG×3 affinity tag after aa 737 (i737) was engineered using the same method as above with oligos (Table S1) targeting the insertion sites and in-frame fused with coding sequence of FLAG×3 at the 5’ end. Fragments containing the amino acid substitutions were moved from pIcsA constructs into pIcsA^737::FLAG^ by restriction cloning with XbaI and HindIII to generate FLAG tagged IcsA substitution mutant constructs.

For extracellular IcsA passenger production, sequences of *icsA* and *icsP* were PCR amplified (Table S1) from *S. flexneri* 2457T chromosomal DNA, digested with NcoI/SalI and NdeI/KpnI, respectively, and then ligated into the MCS1 and MCS2 of the pCDFDuet-1 (Novagen) plasmid sequentially digested in the same way to generate pCDFDuet-1::*icsA-icsP*. A FLAG×3 tag was in-frame inserted after aa 54 (i54) via restriction free cloning (43) with oligos (Table S1) designed to assemble gene blocks containing the coding sequence FLAG×3 tag flanked with sequence up- and down-stream of the i54 via PCR using each other as templates. The gene block was then used as megaprimer pairs to PCR clone the FLAG×3 tag at i54 using pCDFDuet-1::*icsA-icsP* as the template to generate the co-expression construct pCDFDuet-1::*icsA*^*54::FLAG*^*-icsP* (pIcsA-IcsP).

### IcsA passenger purification and size exclusion chromatography

For IcsAp (IcsA^54::FLAG^) production, C43(DE3) co-expressing IcsP and FLAG-tagged IcsA were subcultured 1 in 20 into 4 L of LB from 18 h cultures and were grown to an OD_600_ of 0.4-0.6 at 37 °C. Expression of IcsA was then induced with 1 mM isopropyl β-D-1-thiogalactopyranoside (IPTG) and cultures were incubated at 30 °C for 18 h. The culture supernatant was harvested via centrifugation (7,000 ×g, 25 °C, 15 min), and further cleaned by passing through an asymmetric polyethersulfone (aPES) membrane with 0.2 μm pore size (Rapid-flow, Thermo Scientific). Proteins in the filtered culture supernatant were then concentrated and exchanged into 50 ml of TBS buffer [50 mM Tris, 150 mM NaCl, pH 7.5] using a VivaFlow 200 filtration system (Sartorius) with 30,000 MWCO PES membranes. The concentrated protein preparation was then loaded onto a TBS pre-equilibrated polypropylene column (Thermo Scientific) prepacked with 2 mL of anti-FLAG G1 resin (Genescript). The column was washed with TBS, and IcsA passenger was eluted with 10 ml of 100 mM glycine, pH 3.5. IcsA passenger was then concentrated to 500 μl using a Vivaspin 6 with 10 kDa MWCO (GE Healthcare). For size exclusion chromatography analysis, protein was loaded onto a Superdex 200 Increase 10/300 column (GE Healthcare) with TBS buffer, and different protein elution fractions were pooled and concentrated again as above, which yielded ∼0.8 mg from 4 L culture supernatant. Sizes were only referenced according to the Gel Filtration Makers Kit (for protein molecular weight 29 kDa to 700 kDa, Sigma).

### Polyacrylamide gel electrophoresis and Western immunoblotting

For the analysis of proteins under denatured conditions, bacterial whole cell lysate was prepared by harvesting bacteria (∼5×10 ^8^ cells) grown to the mid-exponential phase (OD_600_∼0.4-0.8) via centrifugation (16,000 ×g, 1 min). The pellet was resuspended into 100 μl SDS-PAGE sample buffer (44) and heated to 98 °C for 10 min. Secreted extracellular protein sample was prepared by harvesting bacterial culture supernatant via centrifugation (5,000 ×g, 5 min) and further clarified through a 0.2 μm filter. Protein was then precipitated using 20% (v/v) trichloroacetic acid at 4 °C, washed by 100% ice-cold acetone and dissolved in 100 μl SDS-PAGE sample buffer in either the presence or absence of DTT (or β-mercaptoethanol (β-ME)) and heated to 100 °C for 5 min. Samples (5-10 μl) were then separated by electrophoresis with either a 12% (v/v) SDS-acrylamide gel or a 4-12% gradient gel (Thermofisher Scientific).

For the separation of the secreted extracellular protein contents in semi-denatured and non-denatured conditions, extracellular protein samples were harvested and clarified as above, and proteins were precipitated with 60% (w/v final) saturated ammonium sulphate at 4 °C overnight. The precipitated extracellular protein was collected via centrifugation (20,000 ×g, 10 min, 4 °C) and dissolved in PBS with 50-fold reduced volume. Sample as above (or affinity purified protein sample) were then mixed 1:1 with the SDS-PAGE sample buffer (semi-denatured condition) or native PAGE sample buffer [62.5 mM Tris pH 6.8, 25% (v/v) glycerol, 1% (w/v) bromophenol blue] (nondenatured condition) in the presence or absence of 10 mM DTT, and unheated protein was separated by electrophoresis with a 4-12% Bis-Tris gel (Thermo Fisher Scientific) with MES running buffer (Thermo Fisher Scientific) (semi-denatured condition); or a hand-casted 8% (v/v) native acrylamide gel [315 mM Tris pH 8.5, 0.1% (w/v) ammonium persulfate (APS), 8% (v/v) acrylamide/bis], with native electrophoresis buffer [25 mM Tris pH 8.5, 192 mM glycine] (nondenatured condition).

Western immunoblotting was done as previously described (11), proteins were detected by rabbit anti-IcsA polyclonal antibodies (pAbs, made in-house) (45), or mouse anti-FLAG antibody (anti-DYKDDDDK antibody, Genscript) with Pierce ECL Western Blotting Substrate (Thermo Fisher Scientific).

### Chemical crosslinking

For chemical crosslinking, affinity purified IcsAp protein was mixed with 0.1 mM DSP (Sigma) in PBS and incubated at room temperature for 30 min. The reaction was quenched by 50 mM Tris (pH 7.0). 10 mM DTT was added to reduce the protein crosslink when required. Samples were mixed with SDS-PAGE sample buffer without β - mercaptoethanol (β-ME) and subjected to SDS-PAGE.

### Cell infection assays

All cell infection assays were described previously (11). Briefly, for cell adhesion assay, confluent HeLa cell monolayers were washed with Dulbecco’s PBS D-PBS and 100 μl of *Shigella* bacteria grown at mid-exponential phase washed in MEM were spinoculated (500 ×g, 5 min) onto the monolayer at multiplicity of infection (MOI) of 100 and incubated for 15 min at 37 °C. Unbound bacteria were removed by three PBS washes, and cell associated bacteria were enumerated by resuspending infected monolayers in 0.1% (v/v) Triton X-100 and serial dilution plating onto agar plates.

For cell invasion assays, confluent HeLa cell monolayer were infected with *Shigella* bacteria as above and incubated for 45 min at 37 °C. Monolayers were washed with D-PBS and extracellular bacteria were eliminated by incubating with MEM containing 50 μg/ml of gentamicin. The viable intracellular bacteria were enumerated as above.

For plaque formation assays, MDCK-2 cells were seeded at low density (5× 10^4^ cells per 24-well) and cultured for 5 days to confluent monolayers. *S. flexneri* strains grown to mid-exponential phase were diluted 1:500 in Hank’s balanced salt solution (HBSS) containing 0.1 M sodium citrate and a 250 μl bacterial suspension (10^6^ cells/ml) was added per MDCK-2 monolayer pre-treated with Hank’s balanced salt solution (HBSS) containing 0.1 M sodium citrate (37 °C, 1 h) and incubated for 1.5 h at 37 °C with 5% CO_2_ with gentle rotation every 15 min. At 1.5 h post infection (pi), a 3 ml of DMEM containing 0.5% SeaKem ME agarose (Lonza), 5% (v/v) FBS and 50 μg/ml of gentamicin was added. At 48 h pi, a 3 ml second overlay containing the same media to the first overlay supplemented with 0.1% (w/v) Neutral Red was added and incubated further 2 h before imaging plaques. Plaque diameter was measured by Image J and plotted. For plaque formation with *∆icsB* strains, after 1.5 h pi, a 3 ml of DMEM containing 5% (v/v) FBS, 100 μg/ml of gentamicin and 50 μg/ml of kanamycin were added. The above media was refreshed daily, and plaques were imaged and measured directly using a light microscope with a CCD camera calibrated with a haemocytometer on day 2 pi.

For intracellular F-actin staining, HeLa cells grown to ∼50% confluency on a coverslip were spinoculated (500 ×g, 5 min) with 200 μl *Shigella* bacteria (10^8^ cells) grown at mid-exponential phase (1:1000 diluted in MEM) and incubated for 1 h at 37 °C with 5% CO_2_. At 1 h pi, extracellular bacteria were inactivated by incubation with 500 μl MEM containing 50 μg/ml gentamicin for another 1 h. Coverslips were then washed three times with PBS and were used in subsequent experiments.

### Immunofluorescence microscopy

For IcsA surface labelling, bacteria (10^8^ cells) grown at mid-exponential phase were harvested via centrifugation (16,000 ×g, 1 min), fixed with 3.7% (w/v) formaldehyde in PBS for 20 min at room temperature. Fixed bacteria were washed with PBS and a 5 μl suspension was then centrifuged onto coverslips pre-coated with 0.01% (w/v) poly-L-lysine (Sigma) in a 24-well tray (16,000 ×g, 1 min). Coverslips were then incubated sequentially with rabbit anti-IcsA pAbs (1:100) or together with mouse anti-FLAG (1:100), then with anti-rabbit Alexa Fluor 488 antibody (Invitrogen, 1:100) or together with anti-mouse Alexa Fluor 647 (Invitrogen, 1:100) in PBS containing 10% (v/v) FBS (Gibco) with PBS washes in between. Coverslips were then mounted with ProLong Diamond Antifade Mountant (Invitrogen), and imaged using a ZEISS Axio Vert.A1 microscope.

For intracellular F-actin tail staining, corresponding samples described as above were washed with D-PBS once and fixed with 3.7% (w/v) formaldehyde in PBS for 20 min at room temperature, quenched with 50 mM NH_4_Cl in PBS for 10 min and permeabilised with 0.1% (v/v) Triton X-100 in PBS for 5 min. Samples were then blocked with 5% FBS in PBS for 20 min before stained with Alexa Flour 594 phalloidin (1:100, Invitrogen) for 1 h and 10 μg/ml DAPI (Invitrogen) for 1 min. Samples were mounted and imaged as above.

### Proteinase accessibility assay

Proteinase accessibility assay was described previously (Brotcke-Zumsteg et al., 2014). Briefly, *Shigella* strains grown overnight were collected (1×10^9^ cells) washed with and resuspended into 1 ml PBS and incubated with 33 nM human neutrophil elastase (hNE, Elastin Products) or proteinase K at 20 μg/ml (NEB) at 37 °C. At indicated time points (see the figure legends), 100 µl samples were taken, mixed with equal volume of SDS-PAGE sample buffer and immediately incubated at 100 °C for 15 min. Alternatively, digestion fractions at the indicated time point were centrifuged (16,000 ×g, 1 min) to give whole bacterial and supernatant fractions, mixed with equal volume of SDS-PAGE sample buffer and immediately incubated at 100 °C for 15 min. Samples were then subjected to SDS - PAGE and Western immunoblotting with anti-IcsA antibody or anti-FLAG antibody.

### Structure prediction and annotation

IcsA structures were predicted using Colab Alpha-fold V2.1.0 structure prediction (28) built in Chimera X1.3 (46) and annotated with Chimera X1.3.

## Data Availability

All data generated or analysed during this study are included in this published article (and its Supplementary Information files).

## Acknowledgement

We thank Prof. Kenneth Beagley for providing mammalian cell lines for this study. This study was supported by grants by the Australian National Health and Medical Research Council (NHMRC GNT1144046), the Clive and Vera Ramaciotti Foundations (2017HIG0119) and a Georgina Sweet Award for Women in Quantitative Biomedical Science to MT. JQ received a Faculty of Science Postgraduate Scholarship from the University of Adelaide. This work made use of a microscopy facility supported by the Ian Porter Foundation. All funders had no role in study design, data collection and analysis, decision to publish, or preparation of the manuscript.

## Author contributions

JQ contributed to project conception and design, conducted experiments, and contributed to data collection, analysis and interpretation; MT supervised the study, contributed to data interpretation and obtained the funding. RM contributed to data interpretation and provided study reagents. JQ drafted the manuscript, RM, YH and RM substaintially revised the manuscript.

## Additional Information

The authors declare no competing interests.

